# Anaesthesia with diethyl ether impairs jasmonate signalling in the carnivorous plant Venus flytrap (*Dionaea muscipula*)

**DOI:** 10.1101/645150

**Authors:** Andrej Pavlovič, Michaela Libiaková, Boris Bokor, Jana Jakšová, Ivan Petřík, Ondřej Novák, František Baluška

## Abstract

General anaesthetics are compounds that induce loss of responsiveness to environmental stimuli in animals and humans. The primary site of general anaesthetic action is the nervous system, where anaesthetics inhibit neuronal transmission. Although plants do not have neurons, they generate electrical signals in response to biotic and abiotic stresses. Here, we investigated the effect of the general volatile anaesthetic diethyl ether on the ability to sense potential prey or herbivore attacks in the carnivorous plant Venus flytrap (*Dionaea muscipula*). We monitored trap movement, electrical signalling, phytohormone accumulation and gene expression in response to the mechanical stimulation of trigger hairs and wounding under diethyl ether treatment. Diethyl ether completely inhibited the generation of action potentials and trap closing reactions, which were easily and rapidly restored when the anaesthetic was removed. Diethyl ether also inhibited the later response: jasmonate (JA) accumulation and expression of JA-responsive genes. However, external application of JA bypassed the inhibited action potentials and restored gene expression under diethyl ether anaesthesia, indicating that downstream reactions from JA are not inhibited. Thus, the Venus flytrap cannot sense prey or a herbivore attack under diethyl ether treatment. This is an intriguing parallel to the effect of anaesthesia on animals and humans.

**Highlight:** Carnivorous plant Venus flytrap (*Dionaea muscipula*) is unresponsive to insect prey or herbivore attack due to impaired electrical and jasmonate signalling under general anaesthesia induced by diethyl ether.

## Introduction

The carnivorous plant Venus flytrap (*Dionaea muscipula*) has evolved modified leaves called traps for prey capture (Gibson *et al*., 2009). The trap consists of two lobes that rapidly close in response to a mechanical stimulus delivered to the trigger hairs protruding from the trap epidermis. Two touches of a trigger hair by an insect prey within 20 seconds generate 2 action potentials (APs) that snap the trap in a fraction of second at room temperature (Escalante-Pérez *et al*., 2011; Volkov, 2019). After rapid closure secures the insect prey, the struggling of the entrapped prey in the closed trap results in the generation of further APs that cease to occur when the prey stops moving (Affolter and Olivo, 1975; Libiaková *et al*., 2014). Third, touch and APs increase cytosolic Ca^2+^ levels in digestive glands, which decay if no further APs are triggered (Escalante-Pérez *et al*., 2011; Hedrich and Neher, 2018). Prey struggling, repeated mechanical stimulation and the generation of hundreds of APs result in the accumulation of phytohormones from the jasmonate group (Escalanté-Pérez *et al*., 2011; Libiaková *et al*., 2014; Pavlovič *et al*., 2017). The binding of the isoleucine conjugate of jasmonic acid (JA-Ile) to the CORONATINE INSENSITIVE1 (COI1) protein as part of a coreceptor complex mediates the ubiquitin-dependent degradation of JASMONATE ZIM-DOMAIN (JAZ) repressors, resulting in the activation of jasmonate-dependent gene expression (Staswick and Tiryaki 2004; Chini et al., 2007; Thines et al., 2007; Sheard et al., 2010). Transcriptional activation leads to the synthesis of transport proteins and digestive enzymes that are secreted into the closed trap cavity (Scherzer *et al*., 2013; 2015; 2017; Libiaková *et al*., 2014; Böhm *et al*., 2016a,b). After the prey movement is stopped by exhaustion or death, chemical stimuli from the prey (e.g., chitin, ammonia) enhance the synthesis of digestive enzymes through jasmonate signalling (Libiaková *et al*., 2014; Paszota *et al*., 2014; Bemm *et al*., 2016).

Several lines of evidence indicate that the generation of electrical signals, jasmonate accumulation and expression of genes encoding digestive enzymes are tightly coupled in the Venus flytrap. First, repeated mechanical stimulation of trigger hairs is sufficient to induce accumulation of the well-known bioactive compound JA-Ile within the first hour in stimulated traps (Pavlovič *et al*., 2017). Just two APs are necessary to induce transcription of *JAZ1* within four hours, and after five APs, *JAZ1* transcripts accumulated to the highest level. More than three APs are necessary to induce significant gene expression of digestive enzymes (e.g., dionain and type I chitinase), and the magnitude of expression is dependent on the number of APs triggered. Gene expression can also be triggered by exogenous application of jasmonic acid (JA), JA-Ile or coronatine without any mechanical stimulus. In contrast, application of the JA perception antagonist coronatine-O-methyloxime (COR-MO), which prevents the COI1-JAZ interaction, blocked gene expression despite triggering 60 APs (Böhm *et al*., 2016a; Bemm *et al*., 2016).

Thus, the sequence of signalling events in Venus flytrap resembles the well-known signalling pathway in response to wounding or herbivore attack in ordinary plants (Maffei *et al*., 2007), supporting the hypothesis that botanical carnivory has evolved from plant-defence mechanisms (Pavlovič and Saganová 2015; Bemm *et al*., 2016). Changes in plasma membrane potential followed by fast electrical signals that may travel through the entire plant from the point of origin are amongst the earliest cellular responses to biotic and abiotic stresses in plants (i.e., wounding or herbivore attack, Maffei *et al*., 2007). A breakthrough study for this issue was that by Wildon *et al*. (1992), who for the first time showed the link between electrical signal propagation and biochemical response in tomato plants. Electrical signals are often followed by changes in intracellular Ca^2+^ concentration and generation of reactive oxygen species (e.g., H_2_O_2_, Maffei *et al*., 2007; Kiep *et al*., 2015; Nguyen *et al*., 2018). A direct link between increased cytosolic Ca^2+^ and activation of JA biosynthesis genes by Ca^2+^/calmodulin-dependent phosphorylation of the JJV repressor complex was recently provided by Yan *et al*. (2018). As a result, increased levels of jasmonates (JA-Ile particularly) trigger the expression of JA-responsive pathogenesis (PR)-related proteins through a COI1-JAZ-dependent pathway (De Geyter *et al*., 2012). Recently, we showed that carnivorous plants are not able to distinguish between mechanical stimulation and wounding because these processes share the same signalling pathway with plant defence mechanisms. Both induce electrical signals, jasmonate accumulation and digestive enzyme synthesis, confirming the link among electrical signal propagation, jasmonate accumulation and the expression of digestive enzymes (Krausko *et al*., 2017; Pavlovič *et al*., 2017). Moreover, the secreted enzymes predominantly belong to pathogenesis-related proteins (PR-proteins), indicating that carnivorous plants have exploited their hydrolytic properties, further emphasizing the similarity between botanical carnivory and plant defence mechanisms (Hatano and Hamada, 2008; 2012; Schulze *et al*., 2012).

Recently, we documented that Venus flytraps, sundew traps, *Mimosa* leaves and pea tendrils lost both autonomous and touch-induced movements after exposure to local and general anaesthetics. Anaesthetics also impeded seed germination and chlorophyll accumulation in cress seedlings, indicating that plants under anaesthesia lose responsiveness to environmental stimuli (Yokawa *et al*., 2018; 2019). General anaesthetics (e.g., diethyl ether) are often defined as compounds that induce a reversible loss of consciousness in humans or loss of righting reflex in animals. Anaesthesia can also be defined as loss of responsiveness to environmental stimuli. Clinical definitions are extended to include the lack of awareness to painful stimuli, which is sufficient to facilitate surgical applications in clinical and veterinary practice (Franks, 2008). The primary site of general anaesthetic action in animals and humans is the central nervous system, where these molecules enhance inhibitory neurotransmission or inhibit excitatory neurotransmission (Zhou *et al*., 2012). Although plants do not have neurons and lack a central nervous system, they are able to generate electrical signals (Fromm and Lautner, 2007; Hedrich *et al*., 2016). Claude Bernard (1878) concluded that volatile anaesthetics not only act on neurons but also affect physiological processes in all cells, and different cells have different susceptibilities to volatile anaesthetics, the neurons being the most sensitive (Grémiaux *et al*., 2014). The electrical signals in plants not only trigger rapid leaf movements in ‘sensitive’ plants, such as *Mimosa pudica* or *D. muscipula*, but also induce physiological processes in ordinary plants (Fromm and Lautner, 2007; Mousavi *et al*., 2013). Interestingly, our recent study showed that inhibition of rapid trap closure in Venus flytrap by the general anaesthetic diethyl ether is caused by inhibition of electrical signalling. There were no toxic impacts of the anaesthetics used, and the effects were fully and rapidly reversible after their removal.

Although carnivorous plants still do not belong to the model group of plants, the signalling events described above indicate that Venus flytrap is a suitable model for studying inducibility and plant responses to external stimuli under anaesthesia due to its rapid trap movement. Considering the tight coupling between electrical signal propagation and jasmonate signalling in carnivorous plants (Böhm *et al*., 2016a; Bemm *et al*., 2016; Krausko *et al*., 2017), we hypothesize that anaesthesia can impair not only rapid trap movement triggered by APs but also the cascade of jasmonate signalling events leading to activation of the digestive process. Our study showed that the Venus flytrap cannot sense potential insect prey or a herbivore attack under anaesthesia due to blocked jasmonate signalling.

## Materials and Methods

### Plant material and culture conditions

The Venus flytrap (*D. muscipula* Ellis.) is native to the subtropical wetlands of North and South Carolina on the East Coast of the USA. Experimental plants were grown under standard glasshouse conditions at the Department of Biophysics of Palacký University in Olomouc (Czech Republic) and the Department of Plant Physiology of Comenius University in Bratislava (Slovakia). Well-drained peat moss in plastic pots placed in a tray filled with distilled water to a depth of 1–2 cm was used as a substrate. Daily temperatures fluctuated between 20 and 35°C; relative air humidity ranged from 50% to 100%; and the maximum daily irradiance reached 1500 μmol m^−2^ s^−1^ photosynthetically active radiation (PAR).

### Experimental setup

The Venus flytrap plants were incubated in 15% diethyl ether for 2 h in a polypropylene bag. This was sufficient to anaesthetize the plants, as we found in our previous study, and the plants did not react to mechanical stimulation by rapid trap closure (Yokawa *et al*., 2018). Thereafter, one group of plants served as a nonstimulated control, and the second group was mechanically stimulated or wounded. For this, a small opening in the polypropylene bag was made. For mechanostimulation, the trigger hairs were mechanically stimulated twice within a short period of time and then 40 times with the tip of a pipette (which had been melted by heat and then hardened at room temperature to avoid a wound response by the sharp tip) every 3 min for 2 h (see Pavlovič *et al*., 2017). In the second experiment, a trap was pierced/wounded with a needle twice within a short period of time and then 40 times every 3 min. In one of the experiments, extracellular measurements of electrical signals were performed on a separate group of plants during stimulation (see below). After 2 hours of stimulation, the plants were removed from the bag to allow the plants to recover from anaesthesia. Immediately, the traps from other groups of plants were sampled for phytohormone analysis. Ten hours later, the traps from the third group of plants were sampled for qPCR. At the same time, the nonstimulated traps under diethyl ether for 4 hours were also harvested. Control plants, bagged nonstimulated plants or identically stimulated plants but in the absence of diethyl ether were also harvested for phytohormone analysis and qPCR at the same time points. For each method, different groups of plants were used because wounding during sampling could activate the jasmonate signalling pathway.

### Extracellular measurements of electrical signals

Venus flytrap incubated in diethyl ether for 2 - 4 h in a polypropylene bag with attached electrodes inside was mechanically stimulated or wounded as described above. Mechanical stimulation or wounding was performed through a small opening in the bag. For recovery, the bag was cut off, and the trigger hair was touched repeatedly every 100 s. Control traps without anaesthetics were also measured. The action potentials were measured on the trap surface inside a Faraday cage with non-polarizable Ag–AgCl surface electrodes (Scanlab Systems, Prague, Czech Republic) fixed with a plastic clip and moistened with a drop of conductive EV gel (Hellada, Prague, Czech Republic) commonly used in electrocardiography. The reference electrode was taped to the side of the plastic pot containing the plant submerged in 1–2 cm of water in a dish beneath the pot. The electrodes were connected to an amplifier [gain 1–1000, noise 2–3 μV, bandwidth (–3 dB) 10^5^ Hz, response time 10 μs, input impedance 10^12^ Ω]. The signals from the amplifier were transferred to an analogue–digital PC data converter (eight analogue inputs, 12-bit converter, ±10 V, PCA-7228AL, supplied by TEDIA, Plzeň, Czech Republic), collected every 6 ms (Hlaváčková *et al*., 2006; Ilík *et al*., 2010).

### Quantification of phytohormone tissue level

Two hours after initiation of mechanical stimulation and wounding under anaesthesia (4 hours etherized), trap tissue samples were collected. Control traps without any stimuli under anaesthesia, as well as control traps stimulated (positive control) and non-stimulated (negative control) without anaesthesia were also harvested. The traps were cut off with scissors and immediately (within 10 seconds) frozen in liquid nitrogen and stored at -80°C until analysis. Ten minutes after diethyl ether removal the remaining traps on plants were mechanically stimulated to be sure that plants were only anaesthetized and not dead (the traps had to close). Quantification of JA, JA-Ile, JA-valine (Ja-Val), *cis*-12-oxo-phytodienoic acid (*cis*-OPDA), 9,10-dihydrojasmonic acid (9,10-DHJA), abscisic acid (ABA), salicylic acid (SA), indole-3-acetic acid (IAA) was performed according to the method described by Floková *et al*. (2014). Briefly, frozen plant material (20 mg) was homogenized and extracted using 1 mL of ice cold 10% MeOH/H_2_O (v/v). A cocktail of stable isotope-labelled standards was added as follows: 5 pmol of [^13^C_6_]IAA, 10 pmol of [^2^H_6_]JA, [^2^H_2_]JA-Ile, and [^2^H_6_]ABA, 20 pmol of [^2^H_4_]SA and [^2^H_5_]OPDA (all from Olchemim Ltd, Czech Republic) per sample to validate the LC-MS/MS method. The extracts were purified using Oasis^®^ HLB columns (30 mg/1 ml, Waters) and hormones were eluted with 80% MeOH. Eluent was evaporated to dryness under a stream of nitrogen. Phytohormone levels were determined by ultra-high performance liquid chromatography-electrospray tandem mass spectrometry (UHPLC-MS/MS) using an Acquity UPLC^®^ I-Class System (Waters, Milford, MA, USA) equipped with an Acquity UPLC CSH^®^ C_18_ column (100 × 2.1 mm; 1.7 µm; Waters) coupled to a triple quadrupole mass spectrometer Xevo^™^ TQ-S MS (Waters MS Technologies, Manchester, UK). Two independent technical measurements were performed on four biological replicates.

### Real-time polymerase chain reaction (qPCR)

To study the induction of gene expression in the trap tissue, two corresponding genes of well-characterized proteins from digestive fluid were chosen: the cysteine protease dionain (Schulze *et al*., 2012; Risør *et al*. 2016) and chitinase I (Paszota *et al*., 2014). To determine the effect of anaesthesia on gene expression, we had to find the time point where the induction of gene expression is high. Therefore, we first collected 100 mg of trap tissue from plants after 0, 2, 6, 12, 24, and 48 hours from initiation of mechanical stimulation, wounding or external application of 2 mM jasmonic acid (JA) under a normal atmosphere (air).

Based on this experiment, we chose the 12-hour time point for mechanostimulation and wounding under anaesthesia. First, the plants were enclosed for 2 hours in polypropylene bags with diethyl ether and then repeatedly mechanically stimulated for 2 hours or wounded as described above. After 4 hours of anaesthesia, the plants were removed from the bag. Ten hours later, 100 mg of trap tissue sample was harvested (Fig. S1A). Control plants without anaesthesia in the air were also mechanically stimulated or wounded (positive controls) or were without any stimulation (in the air and under diethyl ether, negative controls).

To determine the effect of JA under diethyl ether treatment, the plants were again enclosed in polypropylene bags with diethyl ether for 2 hours. We applied 2 mM JA to the trap surface (volume dependent on the size of the trap), and the second group of plants had no JA application but was still under a diethyl ether atmosphere. The same was done in the control air-only plants. After 7 hours, trap samples were collected for qPCR analyses (Fig. S1B). The sampling time was shorter than in the previous experiment because prolonged exposure of plants to diethyl ether caused damage to trap tissue.

To exclude the lethal impact of diethyl ether on plants, the recovery of gene expression was also investigated. After 2 hours in diethyl ether, the plants were removed from the bag and kept for 2 hours in the air. Then, the plants were mechanically stimulated or wounded as described above, and after 10 hours, trap samples were collected for qPCR analyses (Fig. S5A).

Samples were stored at -80°C before gene expression analyses. Total RNA was extracted using a Spectrum Plant Total RNA kit (Sigma–Aldrich, USA), and DNase I was added and purified using an RNA Clean & Concentrator kit (Zymoresearch, USA) according to the manufacturer’s instructions. The RNA integrity was assessed by agarose (1%) gel electrophoresis. The concentration and sample purity were measured by a NanoDrop™ 1000 spectrophotometer (Thermo Fisher Scientific, Germany). The synthesis of the first strand of cDNA was performed by an ImProm-II Reverse Transcription System (Promega) using Oligo(dT)15 primers according to the manufacturer’s protocol. The primers (Tab. S1) for *dionain, chitinase I* and the reference gene *actin* were designated by the Primer3plus tool (http://primer3plus.com/web_3.0.0/primer3web_input.htm). Gradient PCR was used to determine the annealing temperature (T_a_) of the primers (Tab. S1). Each amplified product was assessed by agarose (2%) gel electrophoresis and subsequently sequenced by the Sanger method to verify product specificity. The stability of the reference genes was evaluated by the 2^−ΔCT^ method (Livak and Schmittgen, 2001) and the BestKeeper tool (http://www.gene-quantification.info/). Actin represented a suitable reference gene that was not affected by the different treatments (data not shown). For real-time PCR, specific gene sequences were amplified by Maxima SYBR Green/ROX qPCR Master Mix (Thermo Fisher Scientific). Real-time PCR reactions were performed in 96-well plates on a Light Cycler II 480 (Roche) device, and the relative changes in gene expression were estimated according to Pfaffl (2001). All samples for PCR experiments were analysed in four biological and three technical replicates.

### Western blotting

To detect and quantify cysteine protease (dionain) and type I chitinase, polyclonal antibodies against these proteins were raised in rabbits by Agrisera (Vännäs, Sweden) and Genscript (Piscataway, NJ, USA). The following amino acid sequences (epitopes) were synthesized: cysteine protease, (NH_2_-) CAFQYVVNNQGIDTE (-CONH_2_) (Agrisera, Vännäs, Sweden), and chitinase I, (NH_2_-) CTSHETTGGWATAPD (-CONH_2_) (Genscript, Piscataway, NJ, USA), as we described previously (Pavlovič *et al*., 2017). All sequences were coupled to a carrier protein (keyhole limpet hemocyanin, KLH) and injected into two rabbits each. The terminal cysteine of the peptide was used for conjugation. The rabbit serum was analysed for the presence of antigen-specific antibodies using an ELISA.

The digestive fluid was collected 48 hours after the beginning of mechanical stimulation and wounding from plants in the air (no digestive fluid was secreted under diethyl ether). The samples were heated and denatured for 30 min at 70°C and mixed with modified Laemmli sample buffer to a final concentration of 50 mM Tris-HCl (pH 6.8), 2% SDS, 10% glycerol, 1% β-mercaptoethanol, 12.5 mM EDTA, and 0.02% bromophenol blue. The same volume of digestive fluid was electrophoresed in 10% (v/v) SDS polyacrylamide gel (Schägger, 2006). The proteins in the gels were either visualized by silver staining (ProteoSilver; Sigma Aldrich) or transferred from the gel to a nitrocellulose membrane (Bio-Rad) using a Trans-Blot SD Semi-Dry Electrophoretic Transfer Cell (Bio-Rad). After blocking in TBS-T containing 5% BSA overnight, the membranes were incubated with the primary antibody for 1 h at room temperature, and after washing, the membrane was incubated with the secondary antibody: the goat antirabbit IgG (H + L)-horseradish peroxidase conjugate (Bio-Rad). Blots were visualized by an Amersham Imager 600 gel scanner (GE HealthCare Life Sciences, Tokyo, Japan).

### Statistical analyses

All data are from biological replicates, and each biological sample was analysed in two or three technical replicates. Significant differences between treatments were evaluated by two-tailed Student’s *t*-test (Microsoft Excel).

## Results

### Anaesthesia inhibits electrical signalling and trap closing reactions

The trap of the Venus flytrap plant generates typical APs in response to mechanical stimulation of trigger hair or wounding. Two APs resulted in rapid trap closure within a second (Fig. 1A, Video S1, S4). The shape, duration and amplitude of APs triggered by mechanostimulation and wounding were the same (Fig. S2). However, after 2 hours under diethyl ether anaesthesia, the trap lost the closing response and the ability to generate APs in response to both stimuli (Fig. 1B, Video S2, S5). One hundred seconds after removal of diethyl ether, APs with a reduced amplitude and increased half-width were detected (Fig. 2A), but they were not able to trigger trap closure. In some traps, the first AP was detected after 200 s during recovery. The amplitude and spike half-width of the recorded APs gradually recovered (recorded every 100 s, Fig. 2B, C). When the amplitude of APs was lower and the spike half-width longer during recovery, more touches, and thus more APs, were necessary to induce rapid trap closure. The closing response of the trap was fully restored within 10 - 15 minutes, and again, only two touches were sufficient for trap closure after recovery (Video S3, S6).

**Fig. 1.**
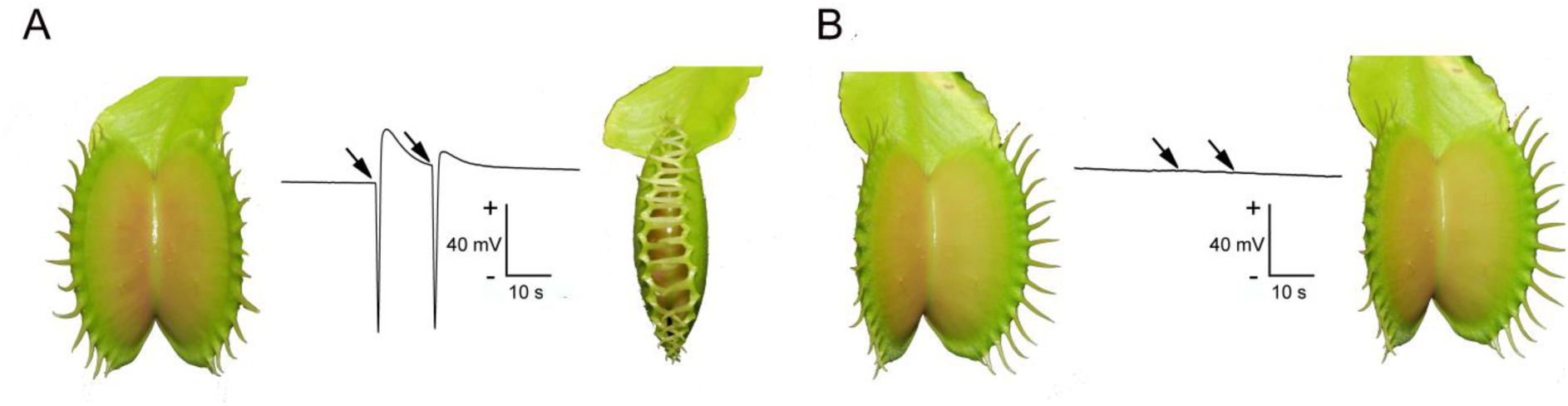
Electrical signalling in the Venus flytrap (*Dionaea muscipula*) under anaesthesia with diethyl ether. (A) Two touches of trigger hairs (arrows) or wounds generate two action potentials and rapid trap closure. (B) Action potentials are not generated in response to two touches (arrows) when exposed to diethyl ether.

**Fig. 2.**
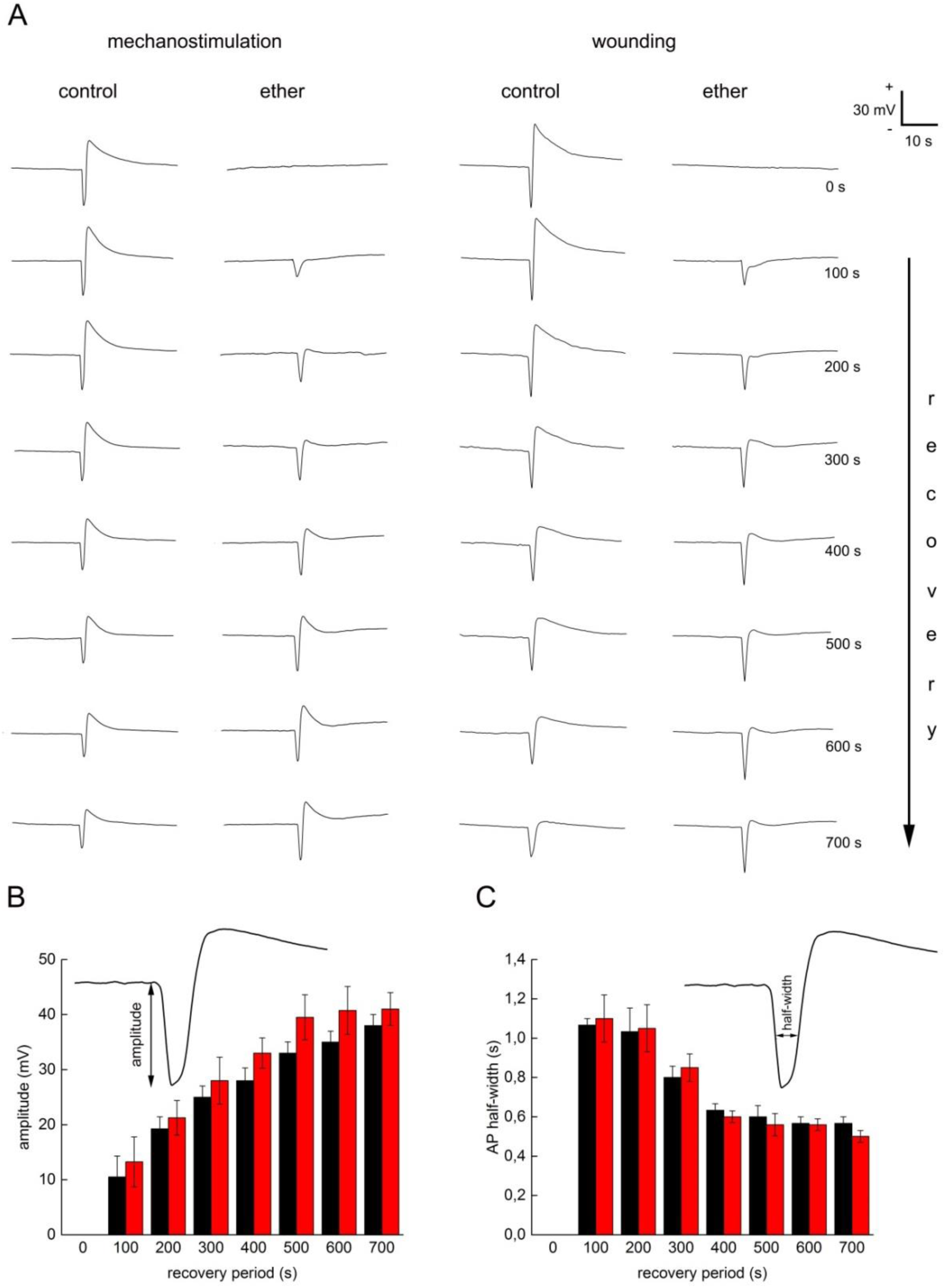
Recovery of electrical signalling after removing of diethyl ether in the Venus flytrap (*Dionaea muscipula*). (A) Recovery of action potentials in response to mechanical stimulation and wounding. The representative APs from four independent experiments are depicted. (B) Recovery of action potential amplitude. (C) Recovery of action potential spike half-width. Black bars – mechanostimulation, red bars – wounding. Means ± S.E., n = 4. There were not significant differences between APs generated in response to wounding and mechanical stimulation.

### Anaesthesia inhibits the accumulation of jasmonates

In our previous study, we found that the jasmonate tissue level in Venus flytrap was the highest within the first two hours of stimulation (Pavlovič *et al*., 2017). Therefore, we chose this time point for phytohormone analysis under anaesthesia. We found a clear activation of the JA signalling pathway for both mechanostimulation and wounding, consistent with our previous study (Pavlovič *et al*., 2017). There was more than a 300-fold increase in the JA tissue level for both types of stimulation in the air (Fig. 3A, I). The bioactive compound JA-Ile increased 23- and 13-fold in response to mechanostimulation and wounding, respectively (Fig. 3B, J). JA-Val significantly increased only in response to wounding (Fig. 3C, K), and the content of *cis*-OPDA did not change significantly (Fig. 3D, L). Under anaesthesia, diethyl ether completely inhibited jasmonate accumulation in response to mechanostimulation (Fig. 3A-E). In response to wounding, there was only a slight but significant increase of jasmonates. The JA level increased 7-fold, and the bioactive JA-Ile level increased 1,5-fold (Fig. 3I, J). This increase probably did not reach the threshold level for activation of the JA signalling pathway, as indicated by the qPCR data (see below). The levels of other plant hormones (ABA, IAA) did not change significantly (Fig. 3F, H, N, P). There was a trend towards a 2-fold increase in SA level in response to both stimuli, irrespective of treatment (air vs. diethyl ether Fig. 3G, O).

**Fig. 3.**
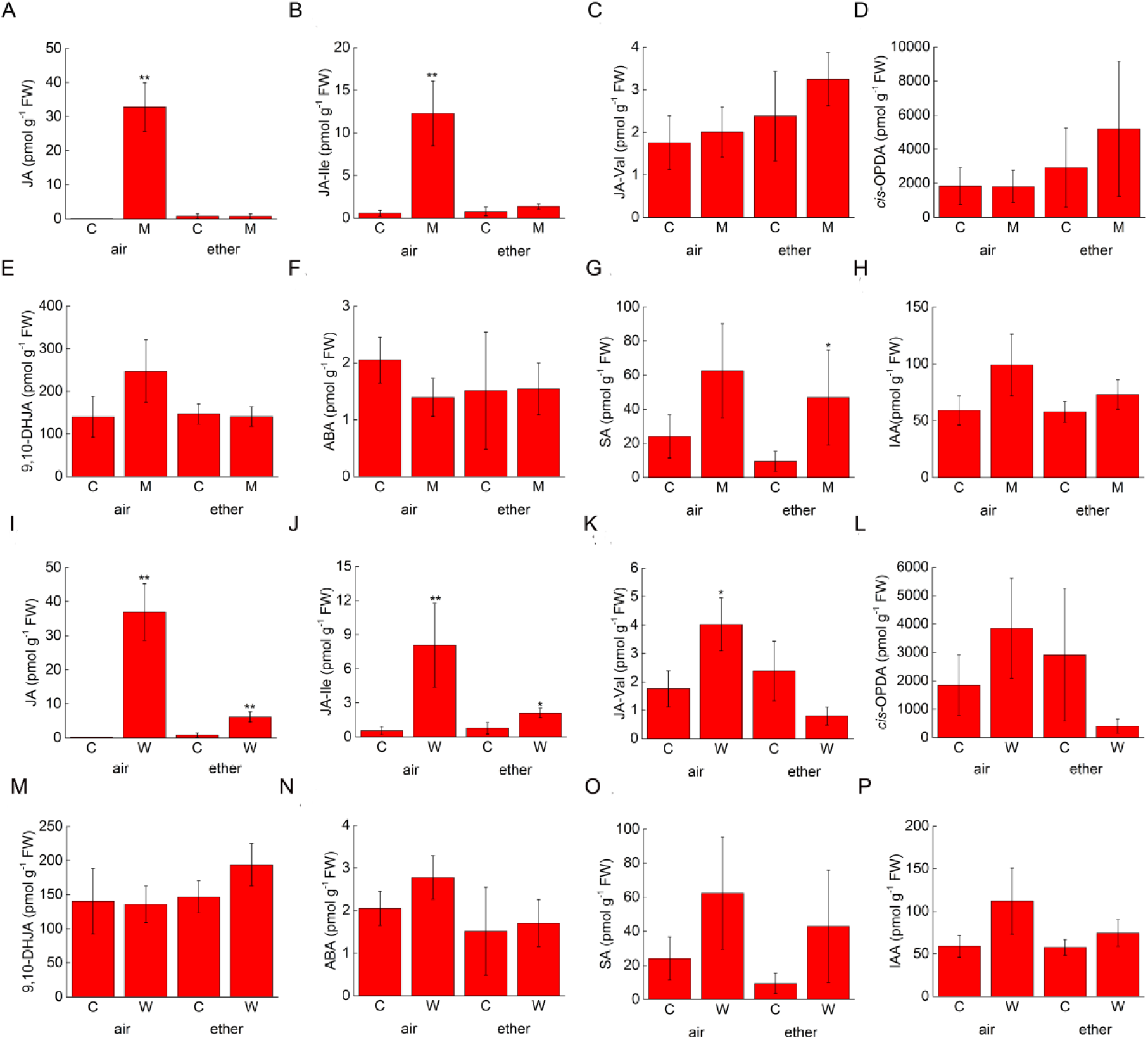
Accumulation of phytohormones in trap tissue of the Venus flytrap (*Dionaea muscipula*) two hours after mechanostimulation and wounding in the air and under anaesthesia with diethyl ether. (A-H) mechanostimulation, (I-P) wounding. (A,I) Jasmonic acid (JA), (B,J) isoleucine conjugate of jasmonic acid (JA-Ile), (C,K) valine conjugate of jasmonic acid (JA-Val), (D,L) *cis*-12-oxo-phytodienoic acid (*cis*-OPDA), (E,M) 9,10-dihydrojasmonic acid (9,10-DHJA), (F,N) abscisic acid (ABA), (G,O) salicylic acid (SA), (H,P) indole-3-acetic acid (IAA). C - control, M -mechanostimulation, W - wounding. Means ± S.D. from four biological replicates, n = 4. Significant differences evaluated by two-tailed Student *t*-test between control and mechanostimulation and control and wounding in the air or diethyl ether are denoted by asterisks at P < 0.01 (**), P < 0.05 (*).

### Transcription of jasmonic acid-responsive genes is inhibited under anaesthesia

First, we analysed the time dependence of the mRNA levels of two selected JA-responsive genes in Venus flytrap: the cysteine protease *dionain* and *chitinase I* (Böhm *et al*., 2016; Bemm *et al*., 2016a). The kinetics of the upregulation of mRNA levels for both genes were similar. The highest mRNA level was found between 12 and 24 hours after the first AP was triggered for both types of stimulation. At 48 hours, the mRNA levels declined. The kinetics of mRNA levels were different for the external application of JA; the mRNA levels of both genes gradually increased over 48 hours (Fig. S3A, B). The protein product of these genes was detected in digestive fluid after 48 hours (Fig. S4). Based on these results, we chose the shortest possible time point, where the upregulation of gene expression was evident, to investigate the effect of anaesthesia. In this experiment, plants were exposed to diethyl ether for two hours, and the traps were mechanostimulated or wounded for the next two hours. The diethyl ether was removed, and the traps were sampled 10 hours later (12 hours after the first mechanostimulus, Fig. S1A). Fig. 4 clearly shows that the mRNA levels of both investigated genes (*dionain, chitinase I*) were not increased under anaesthesia and were comparable with the nonstimulated control in the air or diethyl ether. Two hours after removing diethyl ether, the Venus flytrap was again able to upregulate gene expression in response to mechanostimulation and wounding (Fig. S5B, C).

**Fig. 4.**
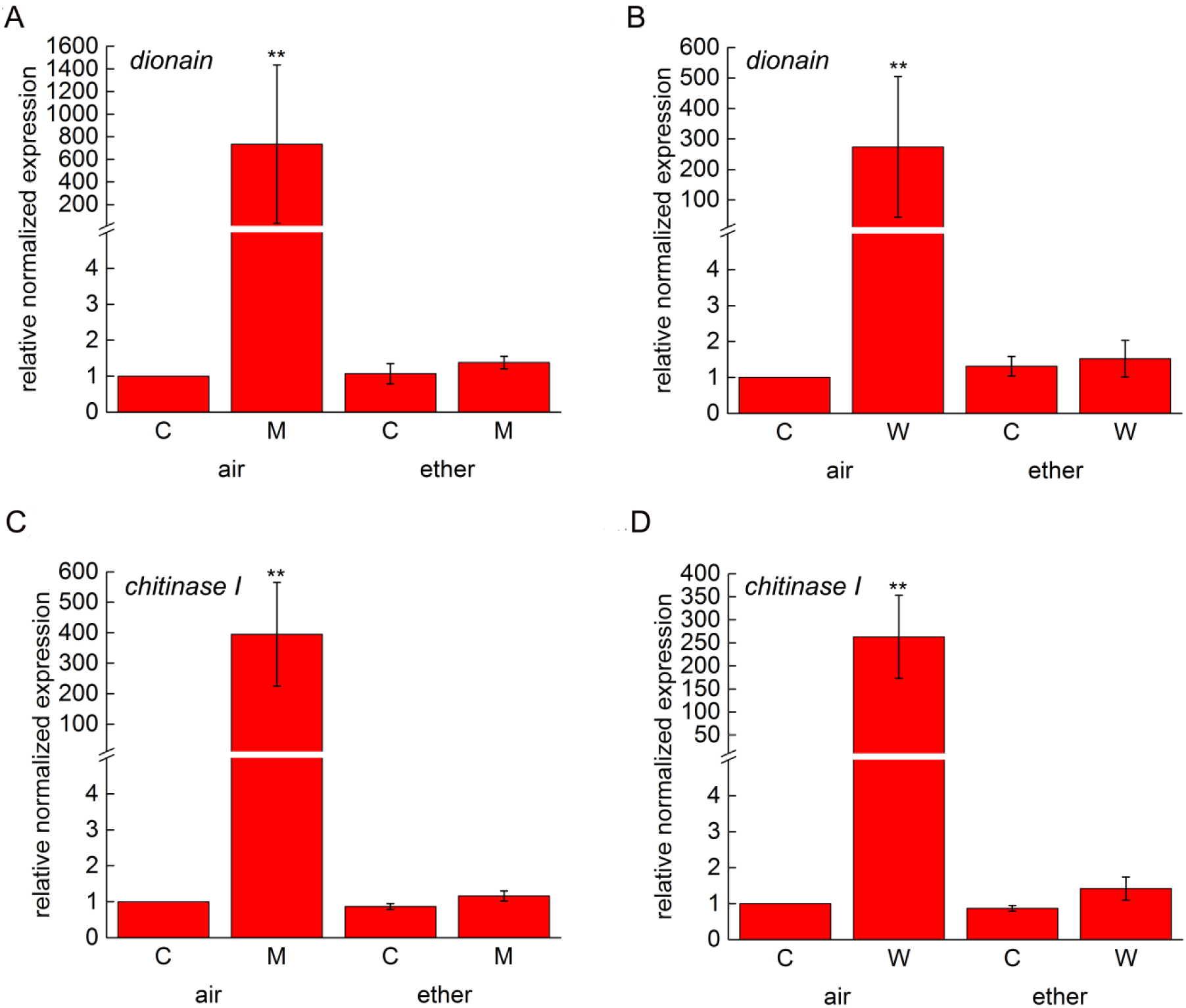
Gene expression in the air and under anaesthesia with diethyl ether in the Venus flytrap (*Dionaea muscipula*). The traps were kept in the air or under diethyl ether for two hours and then 40-times mechanostimulated or pierced/wounded by needle for the next two hours. Then the diethyl ether was removed and trap tissue was sampled for qPCR after 10 hours. (A) Relative expression of dionain after mechanostimulation and (B) wounding. (C) Relative expression of chitinase after mechanostimulation and (D) wounding. Gene expression for non-stimulated control in the air was set up as 1. Mean expression ± S.E. from four biological replicates (n = 4). Significant differences evaluated by two-tailed Student *t*-test between control and mechanostimulation and control and wounding in the air or diethyl ether are denoted by asterisks at P < 0.01 (**).

### External application of jasmonic acid bypassed electrical signalling and restored gene expression under anaesthesia

To determine whether we can bypass the inhibition of electrical signalling by direct application of JA and thus restore gene expression under anaesthesia, we performed the following experiment. The plants were exposed to diethyl ether for two hours. Then, a few drops of 2 mM JA were applied on the trap surface, and the plants were kept for seven hours under anaesthesia, which was the longest possible time to avoid tissue damage. Then, the traps were sampled for qPCR (Fig. S1B). Fig. 5B shows that JA clearly restored the expression of *chitinase I* under diethyl ether. *Dionain* showed a rather weak nonsignificant response (Fig. 5A); however, at the six-hour time point, the increase was not statistically significant in the experiment depicted in Fig. S3A. This result is consistent with the finding that *chitinase I* expression increased earlier (somewhere between 2-6 hours) than *dionain* (between 6-12 hours, Fig. S3) in response to mechanical stimulation, wounding and JA application.

**Fig. 5.**
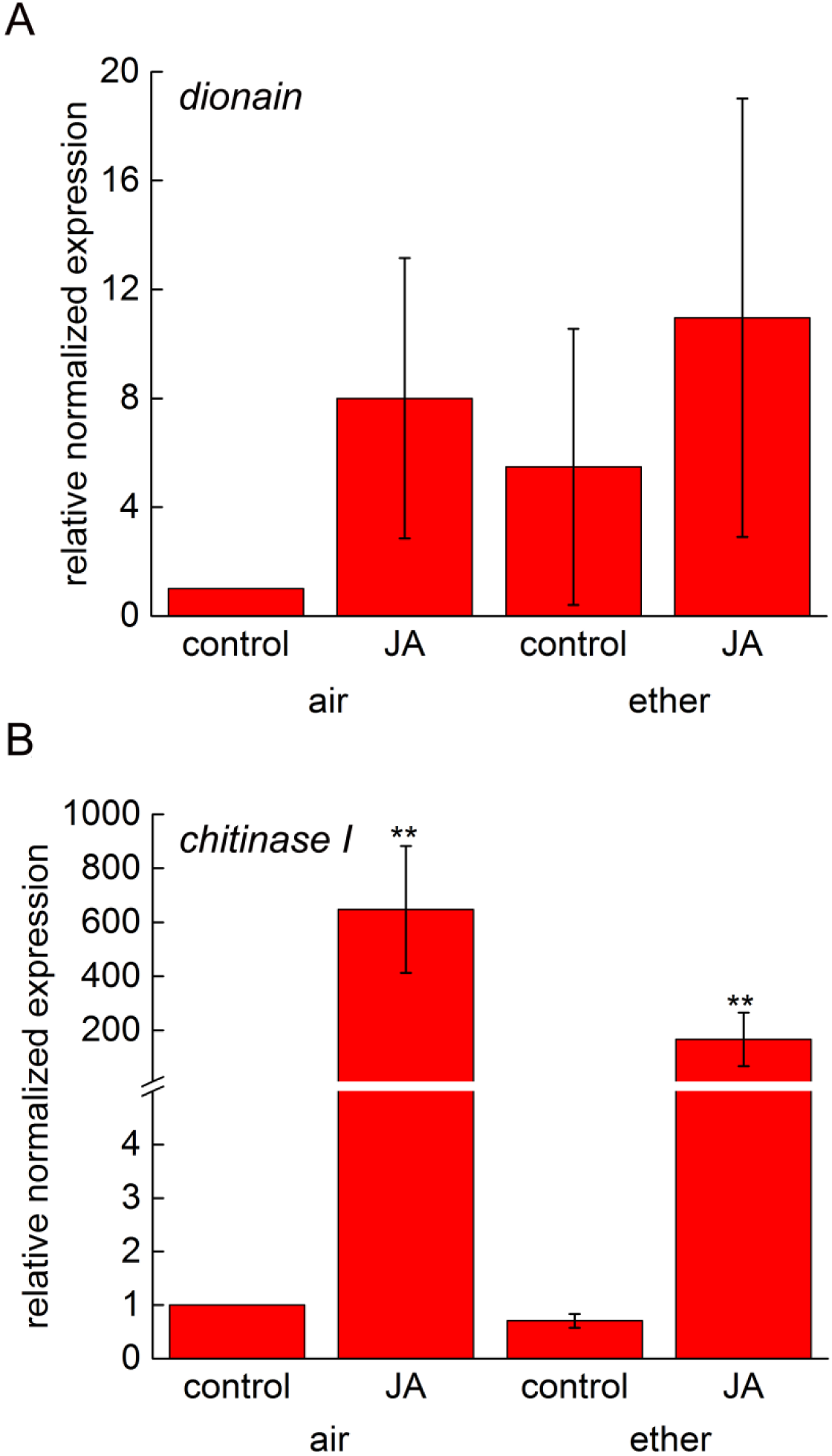
Gene expression in response to exogenous application of 2 mM jasmonic acid after 7 hours in the air or under diethyl ether in the Venus flytrap (*Dionaea muscipula*). The plants were kept in the air or under diethyl ether for two hours and then few drops of jasmonic acid were applied on trap surface. The plants were kept in the same conditions for the next seven hours and then the traps were sampled for qPCR. (A) Relative expression of dionain. (B) Relative expression of chitinase. Mean expression ± S.E. from four biological replicates (n = 4). Significant differences evaluated by two-tailed Student *t*-test between control and mechanostimulation and control and wounding in the air or diethyl ether are denoted by asterisks at P < 0.01 (**).

## Discussion

In this study, we showed that during diethyl ether treatment, Venus flytrap could not sense its environment, and after “waking up”, it did not “remember” what occurred. This was shown by inhibition of APs and trap movement, inability to accumulate jasmonates and no induction of genes encoding digestive enzymes and thus no physiological response. Our observations resemble those in animals and humans where general anaesthesia suppresses central nervous system activity. The volatile anaesthetic halothane (halogenated derivate of ether) produced a concentration-dependent depression of AP amplitude accompanied by an increased spike half-width with complete inhibition at 3 vol % in mammalian nociceptors (MacIver and Tannelian, 1990). Electrical signalling in Venus flytrap was fully recovered in the range of minutes; a similar recovery period was recorded in mammal neurons (MacIver and Tanelian, 1990). During this period, more than two touches were necessary to induce rapid trap closure, supporting the summation of smaller subthreshold charges of APs necessary for trap closing reactions, consistent with electrical memory in Venus flytrap (Volkov *et al*., 2008; 2009). Several previous works indicate that shy plants (*M. pudica*) are also sensitive to anaesthesia. The leaf closing reaction after mechanical stimulus was inhibited by exposure to diethyl ether, halothane and lidocaine but not ketamine (Milne and Beamish, 1999; De Luccia, 2012; Yokawa *et al*., 2018). Although electrical signals were not recorded in these studies, it is tempting to assume that they also inhibited their generation, as electrical signalling and rapid plant movements are tightly coupled (Fromm and Lautner, 2007).

Extensive work has been performed to reveal receptors or mechanisms of anaesthetic action, and two hypotheses have been proposed: the lipid (membrane) theory and protein (receptor) theory (Rinaldi, 2014; Franks, 2008), with several modifications (Lerner *et al*., 1997; Tang and Xu, 2002). Meyer (1899) and Overton (1901) discovered the correlation between the physical properties of general anaesthetic molecules and their potency: the greater the lipid solubility of the compound in olive oil, the greater its anaesthetic potency is. They concluded that solubilization of lipophilic general anaesthetic in the lipid bilayer of the neuron causes its malfunction and anaesthetic effect. Although this simple idea could explain why almost all cells can be anaesthetized, there is also evidence that anaesthetics act by binding directly to sensitive target proteins/receptors. Franks and Lieb (1984) demonstrated that the relationship reported by Meyer (1899) and Overton (1901) could be reproduced using a soluble protein. They showed that a range of general anaesthetics acted as competitive antagonists of the protein firefly luciferase. Remarkably, the inhibition of luciferase was directly correlated with anaesthetic potency, providing persuasive evidence that general anaesthetic drugs could selectively interact with proteins (Weir *et al*., 2006). Until now, many proteins have been shown to contribute to general anaesthesia. Among them are γ-aminobutyric acid type A receptor (GABA_A_), glutamate gated N-methyl-D-aspartate (NMDA) receptors, potassium and sodium channels and others (Mihic *et al*., 1997; Orser *et al*., 2002; Weir, 2006; Zhou *et al*., 2012; Herold and Hemmings, 2012.). Diethyl ether was shown to interact with GABA_A_, NMDA receptors and the potassium channel TREK-1 in animals (Martin *et al*., 1995; Patel *et al*., 1999; Krasowski and Harrison, 2000; Zhou *et al*., 2012). However, the exact nature of general anaesthetic-protein interactions remains a mystery. Anaesthetics may bind to the preformed cavities on proteins by fitting into structurally compatible pockets (key-lock mechanism), causing structural perturbation to the protein channel. Volatile general anaesthetics may have not changed the structure of the membrane channel by a key-lock mechanism but by changing its dynamics by becoming an integral part of amphipathic domains where they can either disrupt the association of the channel with its surroundings or facilitate the formation of structured water clusters within the protein (Tang and Xu, 2002). Another proposed explanation is a combination of the lipid and protein hypotheses: anaesthetics alter the cell membrane properties and may distort the channel protein to block channel function (Lerner *et al*., 1997, Andersen and Koeppe, 2007).

Surprisingly, similar proteins that are suspected as possible targets of volatile anaesthetic diethyl ether in animals and humans have also been discovered in plants, where they are also responsible for electrical signalling. First, glutamate receptor-like proteins (GLRs) in plants are the most closely related proteins to NMDA channels in mammals. They even share similar extensive sequence identity and secondary structure (Lam *et al*., 1998; Weiland *et al*., 2016). GLR3.3 and GLR3.6 are Ca^2+^ channels that mediate the propagation of wound-induced electrical and Ca^2+^ signals in *Arabidopsis* from damaged to undamaged leaves (Mousavi *et al*., 2013; Salvador-Recatalà *et al*., 2014; Hedrich *et al*., 2016; Toyota *et al*., 2018). Moreover, glutamate, which acts as an excitatory neurotransmitter in the vertebrate central nervous system, accumulates in response to wounding in *Arabidopsis*, and GLRs act as sensors that convert the wound signal into an electrical signal that propagates to distant organs where defence responses are induced (Toyota *et al*., 2018). Salvador-Recatalà *et al*. (2014) and Hedrich *et al*. (2016) extended their studies and found that APs triggered by cold water and wounding are not inhibited in local leaves of *glr3.3* and *glr3.6* double mutants. Therefore, the elicitation and propagation of APs is independent of GLR3.3 and GLR3.6 in plants, and they are only important for channelling the signal to neighbouring systemic leaves. This finding is consistent with the results of de Luccia (2012), who found that ketamine, which mediates anaesthesia by blockade of the NMDA receptor in animals, had no effect on the trap closing reaction in Venus flytrap and closing leaflets in *M. pudica*. In addition to Ca^2+^ influx, which is important for the initial depolarization during AP generation, efflux of Cl^−^ accelerates depolarization with the subsequent K^+^ efflux/influx needed for repolarization (Felle and Zimmermann, 2007). The anion QUAC1-type channels and AKT2/3 and SKOR/GORK-type K^+^ channels have bene proposed to be involved in AP generation in plants (van Bel *et al*., 2014; Hedrich *et al*., 2016). Indeed, AKT2 modulates tissue excitability and GORK shapes APs in *Arabidopsis* (Cuin *et al*., 2018). Diethyl ether activated another potassium channel, TREK-1, causing hyperpolarization of the membrane in mammals and inhibiting excitability (Patel *et al*., 1999; Peyronnet *et al*., 2014). The most closely related proteins in plants are TPK potassium channels, which are known to be involved in mechanosensing and controlling membrane potential (Becker *et al*., 2004). The third receptor that was suspected to be a target of diethyl ether anaesthesia in animals is the GABA_A_ receptor. GABA is the main inhibitory neurotransmitter in the central nervous system of vertebrates and exerts its inhibitory effect by activating Cl^−^ currents through the GABA_A_ receptor, hyperpolarizing the membrane and inhibiting excitability. GABA_A_ receptor function is allosterically enhanced by diethyl ether and its halogenated derivatives (Krasowski and Harrison, 2000). Decades ago, GABA was shown to rapidly accumulate in plant tissue in response to different biotic and abiotic stresses, but its receptor was unknown. Recently, the first GABA receptor was found in plants and identified as aluminium-activated malate transporter (ALMT, Ramesh *et al*., 2015; Žárský, 2015). Activation of ALMT results in depolarization of the membrane, and inversely, GABA inhibition results in hyperpolarization of the membrane potentials, generating a state of low excitability (Žárský, 2015). Although the outcome is surprisingly similar to the effect of GABA on animal neurons, there is no sequence homology to the GABA_A_ receptor except for the small region responsible for the GABA interaction (Ramesh *et al*., 2015). This finding decreases the probability that diethyl ether may have the same effect on two unrelated proteins, even if it is only a positive allosteric modulator. As we lack exact data that would allow us to identify the molecular bases underlying the initiation and propagation of APs in plants, it is impossible to identify the protein target of anaesthetic on electrical signals in plants. Either other molecules represent targets of diethyl ether in plants or the membrane theories proposed by Meyer (1899), Overton (1901), and Lerner *et al*. (1997) are relevant in the case of plants.

There is an intriguing parallel to the effects of anaesthetics on animals and humans. Anaesthesia induces loss of responsiveness to environmental stimuli as well as loss of pain perception during surgical operation. Pain sensing in humans results from the action of prostaglandins on peripheral sensory neurons (nociceptors) and on central sites within the spinal cord and the brain (Funk *et al*., 2001; Ricciotti and FitzGerald, 2011). Tissue injury triggers cyclooxygenase-2 (COX-2) in peripheral tissue to convert arachidonic acid to prostaglandin E2 (PGE2), resulting in stimulation/sensitizing of the nociceptor in peripheral nerve to send a signal for pain to the central nervous system. The oxylipin pathway leading to prostaglandin synthesis in animals is mimicked in plants by a similar pathway that leads to the synthesis of jasmonates (Pan *et al*., 1998). We do not claim that plants feel pain, but they use structurally similar molecules as warning signals. We believe that the suppression of jasmonate accumulation under anaesthesia is mediated by the inhibition of electrical signalling, which is tightly coupled to the JA response in ordinary (Mousavi *et al*., 2013; Toyota *et al*., 2018) and carnivorous plants (Böhm *et al*., 2016a; Bemm *et al*., 2016). However, the mechanism of action strongly differs between animals and plants. Whereas the production of oxylipins in plants is mainly downstream from electrical signalling, in animals, it is upstream (prostaglandins sensitize nociceptors for pain). Thus, under anaesthesia, the warning signal (prostaglandins) in animals can be synthesized but is not sensed; in plants, the warning signal (JA) is not synthesized at all. Because of this, exogenous application of JA under anaesthesia can bypass inhibited electrical signalling in plants and trigger the response (Fig. 5).

Wounding, cutting, burning or herbivore attack can all induce electrical signalling and jasmonate accumulation in ordinary plants (Herde *et al*., 1996; 1999; Maffei *et al*., 2007; Mousavi *et al*., 2013). Several genome-wide transcript profiling studies have demonstrated that jasmonates trigger extensive transcriptional reprogramming of metabolism. Jasmonates directly mediate the crucial switch from growth to defence, enabling the plant to reallocate energy to protect itself; thus, plant defence represents a significant cost for plants. JA-induced expression of defence genes occurred concomitantly with the repression of photosynthetic genes and genes involved in cell division and expansion (Stintzi *et al*., 2001; Światek *et al*., 2002; Giri *et al*., 2006; Pauwels *et al*., 2009; Attaran *et al*., 2014). Plants with constitutively activated JA signalling (e.g., *cev1*) exhibit stunted growth (Ellis and Turner, 2001). Additionally, plants with activated JA signalling in an herbivore–free environment have decreased seed production (Baldwin, 1998; Cipollini, 2006). In this study, we showed that we can turn off jasmonate signalling and suppress the production of warning signals and stress by anaesthesia in plants as in animals. Transplantation, wounding, cutting and grafting in horticultural practices are shocks for plants that affect their physiology and production. It is tempting to assume that doing this under anaesthesia can significantly improve plant fitness. This may explain the historical record of successful transplantation of large trees under chloroform treatment into new ground without significant damage by the famous Indian botanist Sir Jagadish Chandra Bose (Yokawa *et al*., 2019). Similar to pain in humans during surgery, anaesthetics inhibit the production of warning signals in plants. Although wounding slightly increased JA and JA-Ile under anaesthesia in this study, this increase probably did not reach the threshold level for activation of gene expression. Because JA synthesis is controlled at the level of substrate availability (Koo and Howe, 2009), this slight increase can be explained by the direct release of lipids from damaged membranes without activation of phospholipases that release JA precursors (e.g., linolenic acid) from plastid lipids (Ishiguro *et al*., 2001; Hyun *et al*., 2008). For phospholipase activation, signalling events (cytoplasmic Ca^2+^ increase) are important (Ryu and Wang 1996; Wang *et al*., 2000). Calcium is involved in the generation of APs in Venus flytrap, and its increased level in the cytoplasm was detected after the third AP (Hodick and Sievers 1988; Krol, 2006; Escalante-Pérez *et al*., 2011), but APs are completely inhibited under anaesthesia. More research is needed on ordinary plants to support these findings, such as how anaesthesia affects the generation of systemic variation potentials (VPs), which are often generated in response to damaging stimuli in ordinary plants.

In conclusion, we showed that the carnivorous plant Venus flytrap cannot sense its environment during anaesthesia with diethyl ether. This situation resembles the effect of general anaesthesia on animals and humans, resulting in a total lack of sensation. After removing the anaesthesia, the recovery of sensitivity is very fast. We have shown that one of the many possible targets of anaesthetics is electrical signals (APs), which affect the later reactions, e.g., generation of warning signals (JA) and transcription of JA-responsive genes. Because jasmonates are important stress hormones that redirect gene expression from growth to defence, the use of anaesthesia during vegetative propagation and plant manipulation in horticultural practice may be plausible, but more experimental studies with ordinary plants are needed. The fact that anaesthesia inhibits electrical signal propagation not only in animals but also in plants and in both affects their sensibility indicates a remarkable similarity between animals and plants.

## Acknowledgements

This work was supported by the Czech Science Foundation Agency [project GAČR 16-07366Y] and Internal Grant Agency of Palacký University [project IGA_PrF_2019_030]. This publication is also the result of the project implementation: Comenius University in Bratislava Science Park supported by the Research and Development Operational Programme funded by the ERDF [Grant number: ITMS 26240220086].

## Author contributions

AP and FB designed the study; AP measured electrical signal, ML and BB performed qPCR, JJ, IP and ON did phytohormone analysis, AP wrote the manuscript and provided materials and financial support. All authors discussed the results and contributed to the manuscript.

## Supplementary data

**Fig. S1** Timeline of experimental setup for diethyl ether treatment.

**Fig. S2** Comparison of action potentials triggered on the same plant by mechanostimulation (black line) and 200 seconds after by wounding (red line).

**Fig. S3** Timecourse of gene expression in response to mechanostimulation, wounding and external application of jasmonic acid during 48 hours in the Venus flytrap (*Dionaea muscipula*).

**Fig. S4** Protein profile and immunodetection of cysteine protease (dionain) and VF-1 chitinase in the digestive fluid of the Venus flytrap (*Dionaea muscipula*).

**Fig. S5** Recovery of gene expression after anaesthesia in the Venus flytrap (*Dionaea muscipula*).

**Table S1** Primer sequences and properties for the Venus flytrap (*Dionaea muscipula*).

**Movie S1** Mechanical stimulation of trigger hairs twice results in rapid trap closure in the Venus flytrap (*Dionaea muscipula*).

**Movie S2** The trap remains open after repeated mechanical stimulation of trigger hairs under anaesthesia with diethyl ether in the Venus flytrap (*Dionaea muscipula*).

**Movie S3** 700 seconds after removing of diethyl ether the trap reaction to mechanostimulation is restored in the Venus flytrap (*Dionaea muscipula*).

**Movie S4** Wounding the trap by needle triggers rapid trap closure in the Venus flytrap (*Dionaea muscipula*).

**Movie S5** The trap remains open after wounding under anaesthesia with diethyl ether in the Venus flytrap (*Dionaea muscipula*).

**Movie S6** 700 seconds after removing of diethyl ether the trap reaction to wounding is restored in the Venus flytrap (*Dionaea muscipula*).

