## Supplementary data for "Anaesthesia with diethyl ether impairs jasmonate signalling in the carnivorous plant Venus flytrap (*Dionaea muscipula*)"

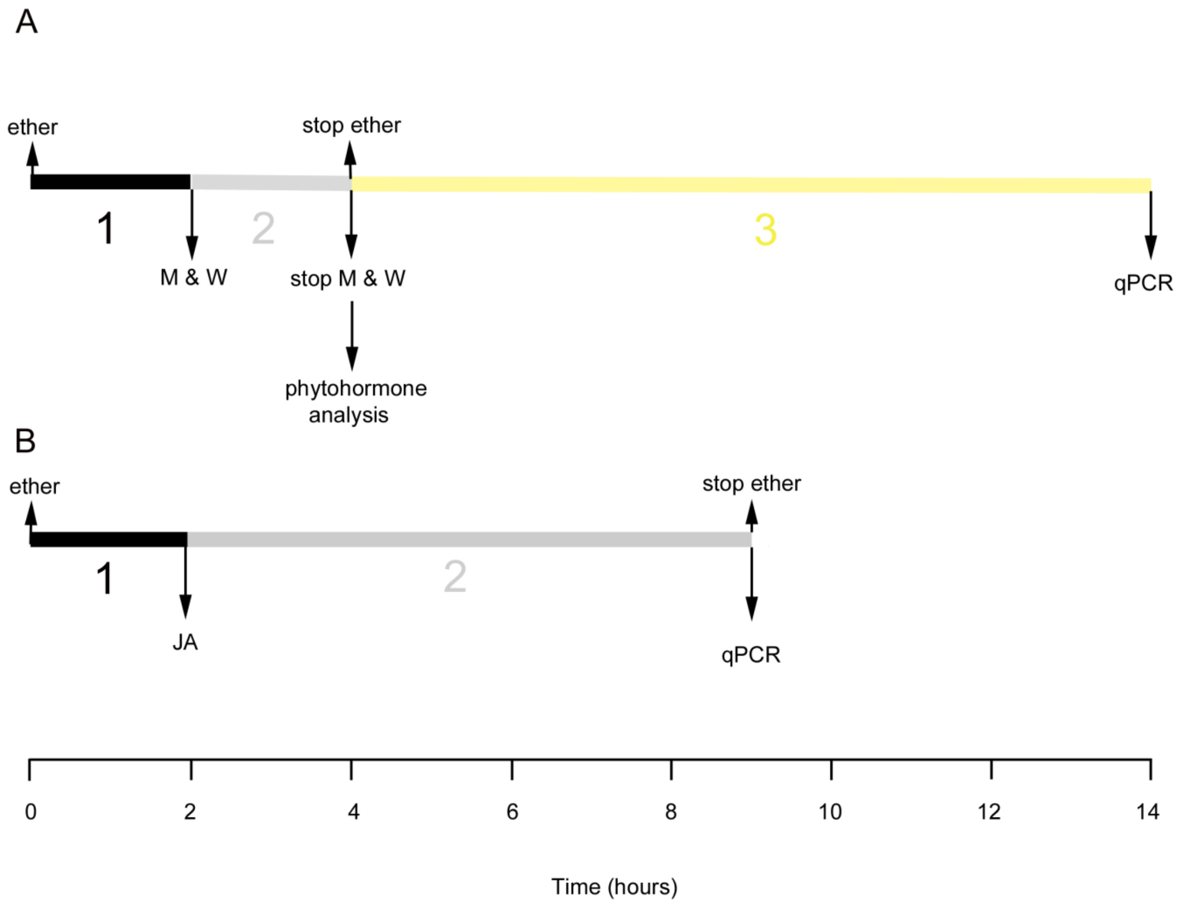


**Fig. S1** Timeline of experimental setup for diethyl ether treatment. (A) The plants were anaesthetized for 2 hours with diethyl ether (black area, 1). Than the trap was mechanically stimulated or wounded (M & W) for the next two hours (grey area, 2). After 4 hours under anaesthesia including 2 hours of stimulation, the trap tissue was sampled for phytohormone analysis. Diethyl ether was removed and after 10 hours trap tissue for qPCR was sampled (yellow area, 3). **(B)** The plants were anaesthetized for 2 hours with diethyl ether (black area, 1). Then few drops of 2mM JA was applied on trap surface for the next seven hours (grey area, 2). Then the trap tissue was sampled for qPCR. For controls black (1) and grey (2) area is in the air. The non-stimulated control under diethyl ether was without any stimuli.

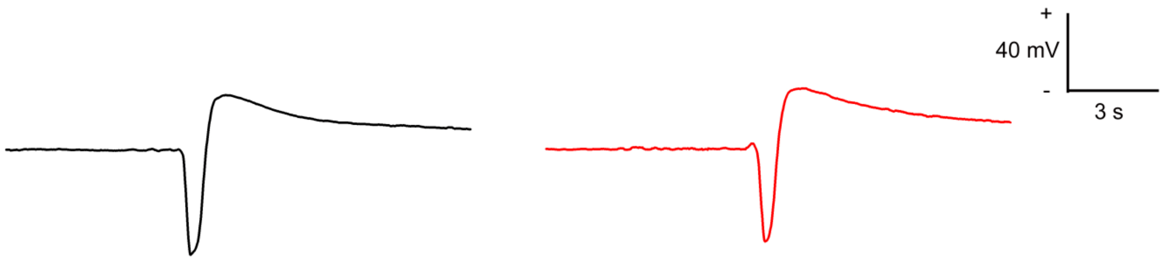


**Fig. S2** Comparison of action potentials triggered on the same plant by mechanostimulation (black line) and 200 seconds after by wounding (red line). There are no significant differences between amplitude, half-width and overall shape of action potential triggered by different stimuli.

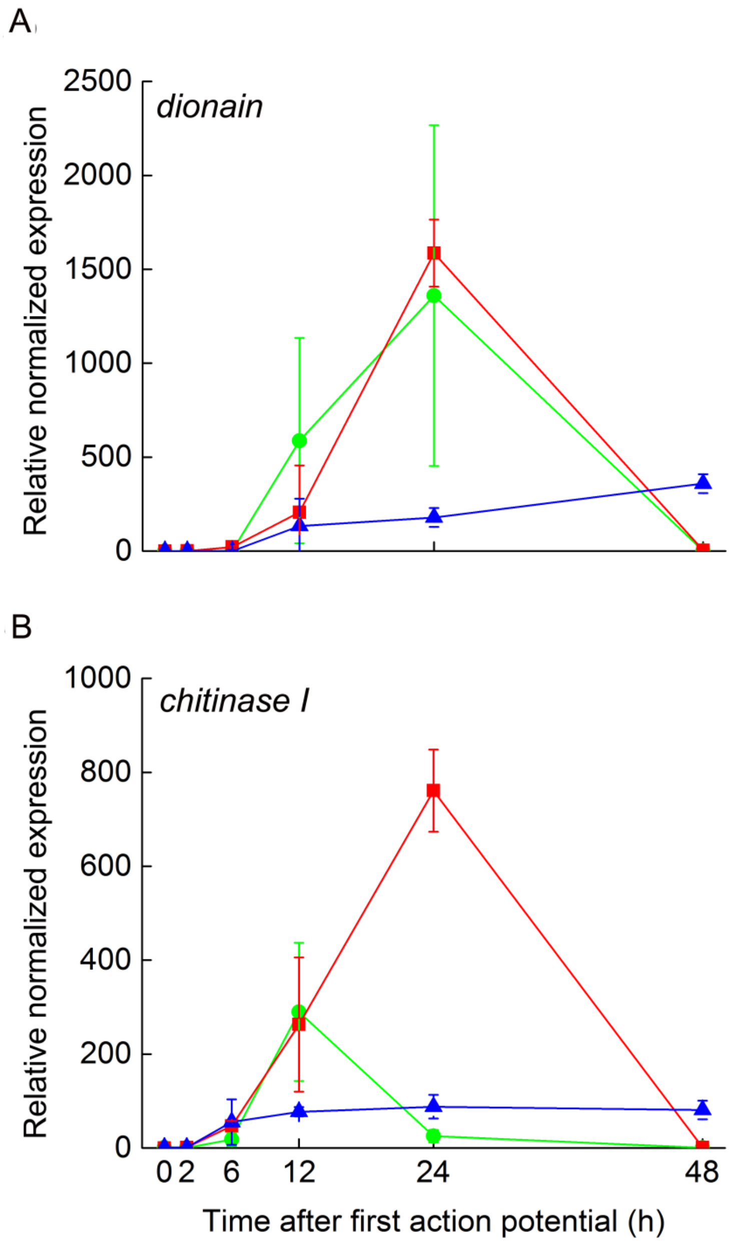


**Fig. S3** Timecourse of gene expression in response to mechanostimulation, wounding and external application of jasmonic acid during 48 hours in the Venus flytrap (*Dionaea muscipula*). The traps were 40-times mechanostimulated or pierced/wounded be needle within the first two hours. Jasmonic acid was applied during whole 48 hours. (A) Cysteine protease dionain, (B) type I chitinase. Mechanostimulation (green circle), wounding (red square), jasmonic acid (blue triangle). Gene expression for 0h (before any stimulation) was set as 1. Mean expression ± S.E. from four biological replicates (n = 4).

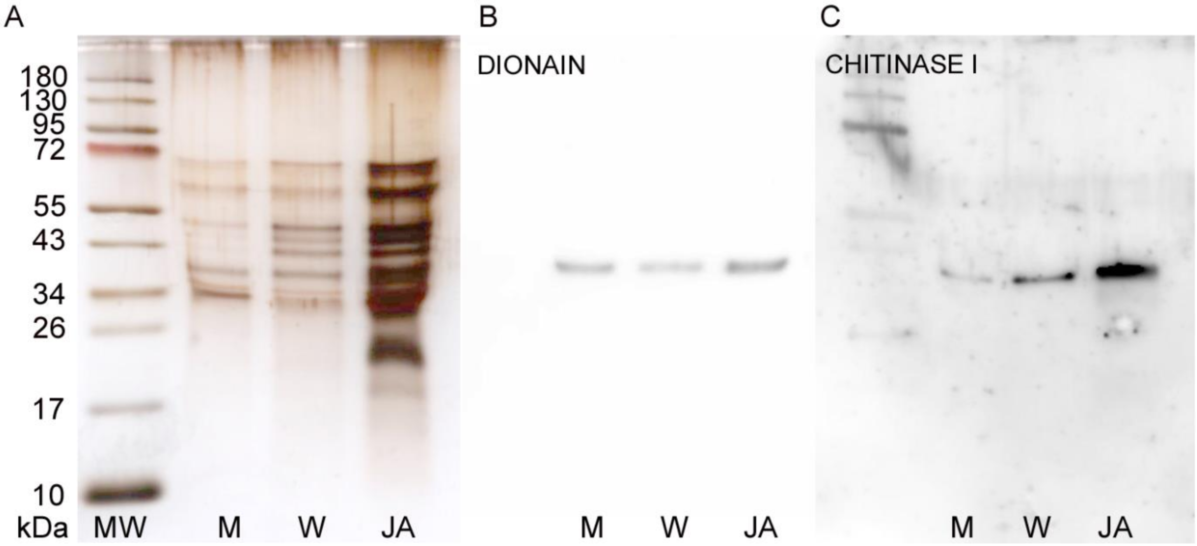


**Fig. S4** Protein profile and immunodetection of cysteine protease (dionain) and VF-1 chitinase in the digestive fluid of the Venus flytrap (*Dionaea muscipula*) after 48 hours. The digestive fluid was collected 48 h after induction, and the proteins were separated in 10% (v/v) SDS polyacrylamide gel and subjected to Western blot analysis. (A) Silver-stained SDS-PAGE of the digestive fluid in response to different stimuli. (B) Western blot analysis of cysteine protease dionain using a protein-specific antibody. (C) Western blot analysis of type I chitinase using a protein-specific antibody. MW, molecular weight; M, mechanical stimulation; W, wounding; JA, 2mM jasmonic acid. The same volume (8 µl) of secreted digestive fluid was loaded. The blots shown are representative of three independent experiments.

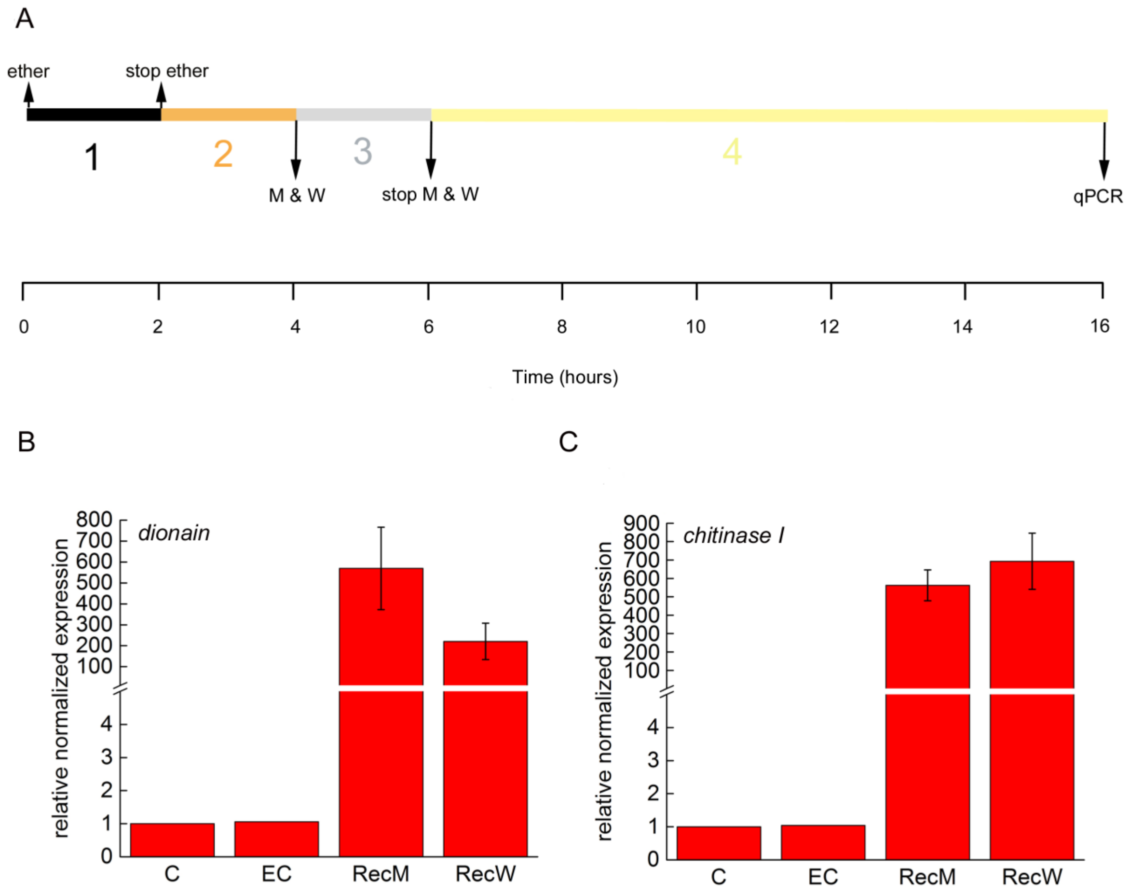


**Fig. S5** Recovery of gene expression after anaesthesia in the Venus flytrap (*Dionaea muscipula*). (A) Timeline of experimental setup for recovery experiments**.** The plants were anaesthetized for 2 hours with diethyl ether (black area, 1). For recovery, the diethyl ether was removed and plants were kept 2 hours in the air (orange area, 2). Than the traps were mechanically stimulated or wounded (M & W) for the next two hours (grey area, 3). After 10 hours, the trap tissue was sampled for qPCR (yellow area, 4). (B) Relative expression of cysteine protease dionain. (C) Relative expression of type I chitinase. Mean expression ± S.E., n = 4. C – control in the air, EC – control in diethyl ether, RecM – recovery after mechanostimulation, RecW – recovery after wounding.

**Table S1 Primer sequences and properties for the Venus flytrap (*Dionaea muscipula*). Ta – annealing temperature.**

| **Primer / accession number** | **Product size (bp)** | **Primer sequence (5´- 3´ direction)** | **$T_a$ (°C)** |
|---|---|---|---|
| ***actin* / KC285589.1.** | 212 | Forward: TCTTTGATTGGGATGGAAGC<br>Reverse: CTCTCTGGAGGAGCAACCAC | 58 |
| ***dionain I* / KP663370.1** | 161 | Forward: ATGGGGCGATGATGATCTTA<br>Reverse: CTTCTCCGCATCATCCTTGT | 58 |
| ***chitinase I* / KF597524.1** | 207 | Forward: GCTCTGCATTTCACCGTTTT<br>Reverse: ACAATGGAGCCAACATCACC | 58 |

**Movie S1** Mechanical stimulation of trigger hairs twice results in rapid trap closure in the Venus flytrap (*Dionaea muscipula*).

**Movie S2** The trap remains open after repeated mechanical stimulation of trigger hairs under anaesthesia with diethyl ether in the Venus flytrap (*Dionaea muscipula*).

**Movie S3** 700 seconds after removing of diethyl ether the trap reaction to mechanostimulation is restored in the Venus flytrap (*Dionaea muscipula*).

**Movie S4** Wounding the trap by needle triggers rapid trap closure in the Venus flytrap (*Dionaea muscipula*).

**Movie S5** The trap remains open after wounding under anaesthesia with diethyl ether in the Venus flytrap (*Dionaea muscipula*).

**Movie S6** 700 seconds after removing of diethyl ether the trap reaction to wounding is restored in the Venus flytrap (*Dionaea muscipula*).
